# OptiFit: an improved method for fitting amplicon sequences to existing OTUs

**DOI:** 10.1101/2021.11.09.468000

**Authors:** Kelly L. Sovacool, Sarah L. Westcott, M. Brodie Mumphrey, Gabrielle A. Dotson, Patrick D. Schloss

## Abstract

Assigning amplicon sequences to operational taxonomic units (OTUs) is often an important step in characterizing the composition of microbial communities across large datasets. OptiClust, a *de novo* OTU clustering method, has been shown to produce higher quality OTU assignments than other methods and at comparable or faster speeds. A notable difference between *de novo* clustering and database-dependent reference clustering methods is that OTU assignments from *de novo* methods may change when new sequences are added to a dataset. However, in some cases one may wish to incorporate new samples into a previously clustered dataset without performing clustering again on all sequences, such as when comparing across datasets or deploying machine learning models where OTUs are features. Existing reference-based clustering methods produce consistent OTUs, but they only consider the similarity of each query sequence to a single reference sequence in an OTU, thus resulting in OTU assignments that are significantly worse than those generated by *de novo* methods. To provide an efficient and robust method to fit amplicon sequence data to existing OTUs, we developed the OptiFit algorithm. Inspired by OptiClust, OptiFit considers the similarity of all pairs of reference and query sequences in an OTU to produce OTUs of the best possible quality. We tested OptiFit using four microbiome datasets with two different strategies: by clustering to an external reference database or by splitting the dataset into a reference and query set and clustering the query sequences to the reference set after clustering it using OptiClust. The result is an improved implementation of closed and open-reference clustering. OptiFit produces OTUs of similar quality as OptiClust and at faster speeds when using the split dataset strategy, although the OTU quality and processing speed depends on the database chosen when using the external database strategy. OptiFit provides a suitable option for users who require consistent OTU assignments at the same quality afforded by *de novo* clustering methods.

**Importance:** Advancements in DNA sequencing technology have allowed researchers to affordably generate millions of sequence reads from microorganisms in diverse environments. Efficient and robust software tools are needed to assign microbial sequences into taxonomic groups for characterization and comparison of communities. The OptiClust algorithm produces high quality groups by comparing sequences to each other, but the assignments can change when new sequences are added to a dataset, making it difficult to compare different studies. Other approaches assign sequences to groups by comparing them to sequences in a reference database to produce consistent assignments, but the quality of the groups produced is reduced compared to OptiClust. We developed OptiFit, a new reference-based algorithm that produces consistent yet high quality assignments like OptiClust. OptiFit allows researchers to compare microbial communities across different studies or add new data to existing studies without sacrificing the quality of the group assignments.

## Introduction

Amplicon sequencing is a mainstay of microbial ecology. Researchers can affordably generate millions of sequences to characterize the composition of hundreds of samples from microbial communities without the need for culturing. In many analysis pipelines, 16S rRNA gene sequences are assigned to operational taxonomic units (OTUs) to facilitate comparison of taxonomic composition between communities to avoid the need for taxonomic classification. A distance threshold of 3% (or sequence similarity of 97%) is commonly used to cluster sequences into OTUs based on pairwise comparisons of the sequences within the dataset. The method chosen for clustering affects the quality of OTU assignments and thus may impact downstream analyses of community composition (1–3).

There are two main categories of OTU clustering algorithms: *de novo* and reference-based. OptiClust is a *de novo* clustering algorithm which uses the distance score between all pairs of sequences in the dataset to cluster them into OTUs by maximizing the Matthews Correlation Coefficient (MCC) (1). This approach takes into account the distances between all pairs of sequences when assigning query sequences to OTUs, in contrast to other *de novo* methods such as the greedy clustering algorithms implemented in USEARCH and VSEARCH (4, 5). In methods employing greedy clustering algorithms, only the distance between each sequence and a representative centroid sequence in the OTU is considered while clustering. As a result, distances between pairs of sequences in the same OTU are frequently larger than the specified threshold, i.e. they are false positives. In contrast, the OptiClust algorithm takes into account the distance between all pairs of sequences when considering how to cluster sequences into OTUs and is thus less willing to take on false positives. A limitation of *de novo* clustering is that different OTU assignments will be produced when new sequences are added to a dataset, making it difficult to use *de novo* clustering to compare OTUs between different studies. Furthermore, since *de novo* clustering requires calculating and comparing distances between all sequences in a dataset, the execution time can be slow and memory requirements can be prohibitive for very large datasets. Reference clustering attempts to overcome the limitations of *de novo* clustering methods by using a representative set of sequences from a database, with each reference sequence seeding an OTU. Commonly, the Greengenes set of representative full length sequences clustered at 97% similarity is used as the reference with VSEARCH (5–7). Query sequences are then clustered into OTUs based on their similarity to the reference sequences. Any query sequences that are not within the distance threshold to any of the reference sequences are either thrown out (closed reference clustering) or clustered *de novo* to create additional OTUs (open reference clustering). While reference-based clustering is generally fast, it is limited by the diversity of the reference database. Novel sequences in the sample will be lost in closed reference mode if they are not represented by a similar sequence in the database. Previous studies found that the OptiClust *de novo* clustering algorithm created the highest quality OTU assignments of all clustering methods (1).

To overcome the limitations of current reference-based and *de novo* clustering algorithms while maintaining OTU quality, we developed OptiFit, a reference-based clustering algorithm. While other tools represent reference OTUs with a single sequence, OptiFit uses multiple sequences in existing OTUs as the reference and fits new sequences to those reference OTUs. In contrast to other tools, OptiFit considers all pairwise distance scores between reference and query sequences when assigning sequences to OTUs in order to produce OTUs of the highest possible quality. Here, we tested the OptiFit algorithm with the reference as a public database (e.g. Greengenes) or *de novo* OTUs generated using a reference set from the full dataset and compared the performance to existing tools. To evaluate the OptiFit algorithm and compare to existing methods, we used four published datasets isolated from soil (8), marine (9), mouse gut (10), and human gut (11) samples. OptiFit is available within the mothur software program.

## Results

### The OptiFit algorithm

OptiFit leverages the method employed by OptiClust of iteratively assigning sequences to OTUs to produce the highest quality OTUs possible, and extends this method for reference-based clustering. OptiClust first seeds each sequence into its own OTU as a singleton. Then for each sequence, OptiClust considers whether the sequence should move to a different OTU or remain in its current OTU, choosing the option that results in a better Matthews correlation coefficient (MCC) (1). The MCC uses all values from a confusion matrix and ranges from negative one to one, with a score of one occurring when all sequence pairs are true positives and true negatives and a score of negative one occurring when all pairs are false positives and false negatives. Sequence pairs that are similar to each other (i.e. within the distance threshold) are counted as true positives if they are clustered into the same OTU, and false negatives if they are not in the the same OTU. Sequence pairs that are not similar to each other are true negatives if they are not clustered into the same OTU, and false positives if they are not in the same OTU. OptiClust iterations continue until the MCC stabilizes or until a maximum number of iterations is reached. This process produces *de novo* OTU assignments with the most optimal MCC given the input sequences.

OptiFit begins where OptiClust ends, starting with a list of reference OTUs and their sequences, a list of query sequences to cluster to the reference OTUs, and the sequence pairs that are within the distance threshold (e.g. 0.03) (Figure 1). Initially, all query sequences are placed into separate OTUs. Then, the algorithm iteratively reassigns the query sequences to the reference OTUs to optimize the MCC. Alternatively, a sequence will remain unassigned if the MCC value is maximized when the sequence is a singleton rather than clustered into a reference OTU. All query and reference sequence pairs are considered when calculating the MCC. This process is repeated until the MCC changes by no more than 0.0001 (default) or until a maximum number of iterations is reached (default: 100). In the closed reference mode, any query sequences that cannot be clustered into reference OTUs are discarded, and the results only contain OTUs that exist in the original reference. In the open reference mode, unassigned query sequences are clustered *de novo* using OptiClust to generate new OTUs. The final MCC is reported with the best OTU assignments. There are two strategies for generating OTUs with OptiFit: 1) cluster the query sequences to reference OTUs generated by *de novo* clustering an independent database, or 2) split the dataset into a reference and query fraction, cluster the reference sequences *de novo*, then cluster the query sequences to the reference OTUs.

**Figure 1:**
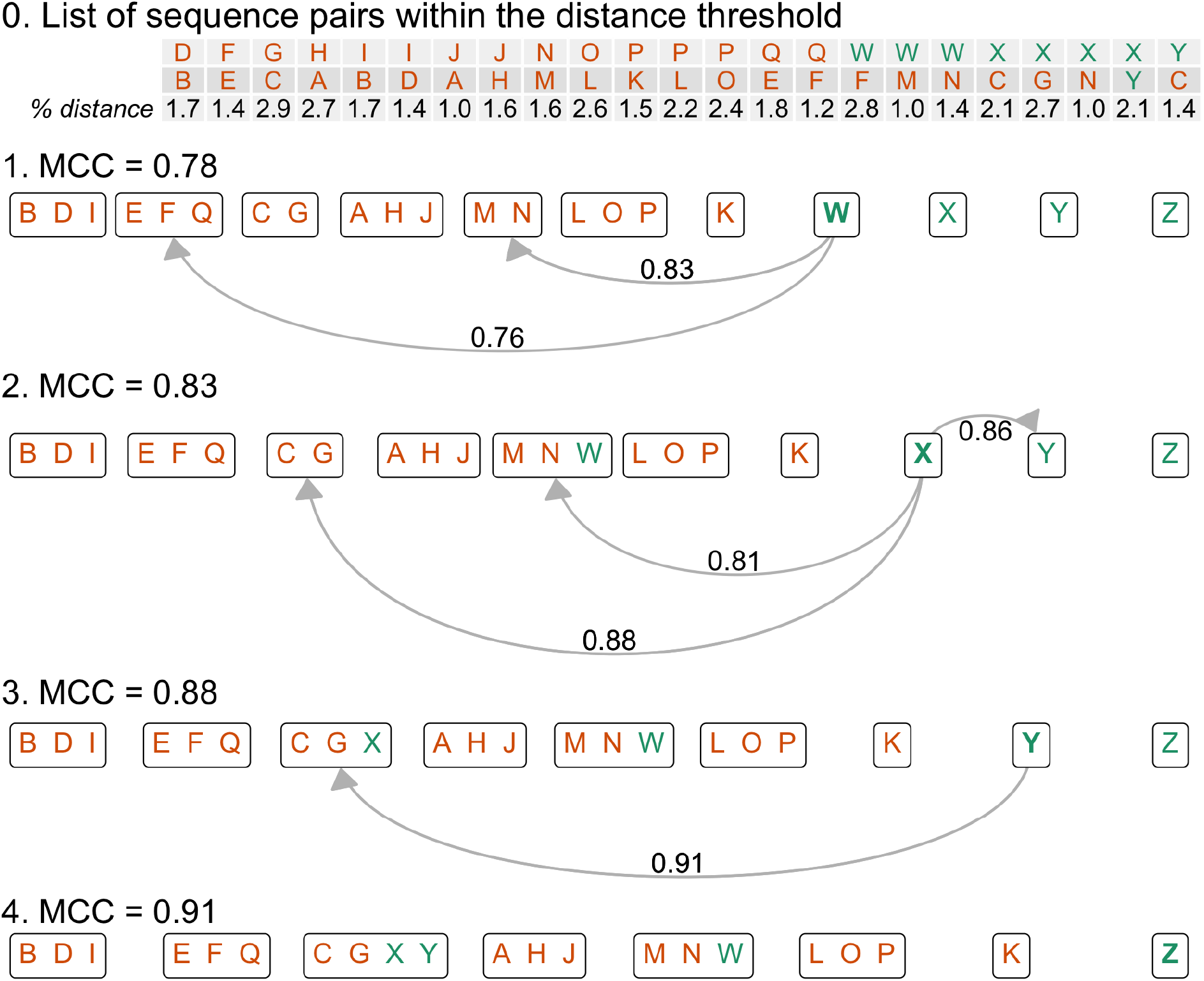
The OptiFit Algorithm. Here we present a toy example of the OptiFit algorithm fitting query sequences to existing OTUs, given the list of all sequence pairs that are within the distance threshold (here 3% is used). The goal of OptiFit is to assign the query sequences W through Z (colored green) to the reference OTUs created by clustering Sequences A through Q (colored orange) which were previously clustered *de novo* with OptiClust (see the OptiClust supplemental text (1)). Initially, OptiFit places each query sequence in its own OTU. Then, for each query sequence (**bolded**), OptiFit determines what the new MCC score would be if that sequence were moved to one of the OTUs containing at least one other similar sequence. The sequence is then moved to the OTU which would result in the best MCC score. OptiFit stops iterating over sequences once the MCC score stabilizes (in this example; only one iteration over each sequence is needed).

### Reference clustering with public databases

To test how OptiFit performs for reference-based clustering, we clustered each dataset to three databases of reference OTUs: the Greengenes database, the SILVA non-redundant database, and the Ribosomal Database Project (RDP) (6, 12, 13). Reference OTUs for each database were created by performing *de novo* clustering with OptiClust at a distance threshold of 3% using the V4 region of each sequence (see Figure 2). After trimming to the V4 region, the databases contained 174,979, 16,192, and 173,648 unique sequences and produced *de novo* MCC scores of 0.72, 0.74, and 0.73 for Greengenes, RDP, and SILVA, respectively. Clustering sequences to Greengenes and SILVA in closed reference mode performed similarly, with median MCC scores of 0.85 and 0.77 respectively, while the median MCC was 0.35 when clustering to RDP (Figure 3). For comparison, clustering datasets with OptiClust produced an average MCC score of 0.87. This gap in OTU quality mostly disappeared when clustering in open reference mode, which produced median MCCs of 0.86 with Greengenes, 0.85 with SILVA, and 0.86 with the RDP. Thus, open reference OptiFit produced OTUs of very similar quality as *de novo* clustering, and closed reference OptiFit followed closely behind as long as a suitable reference database was chosen.

**Figure 2:**
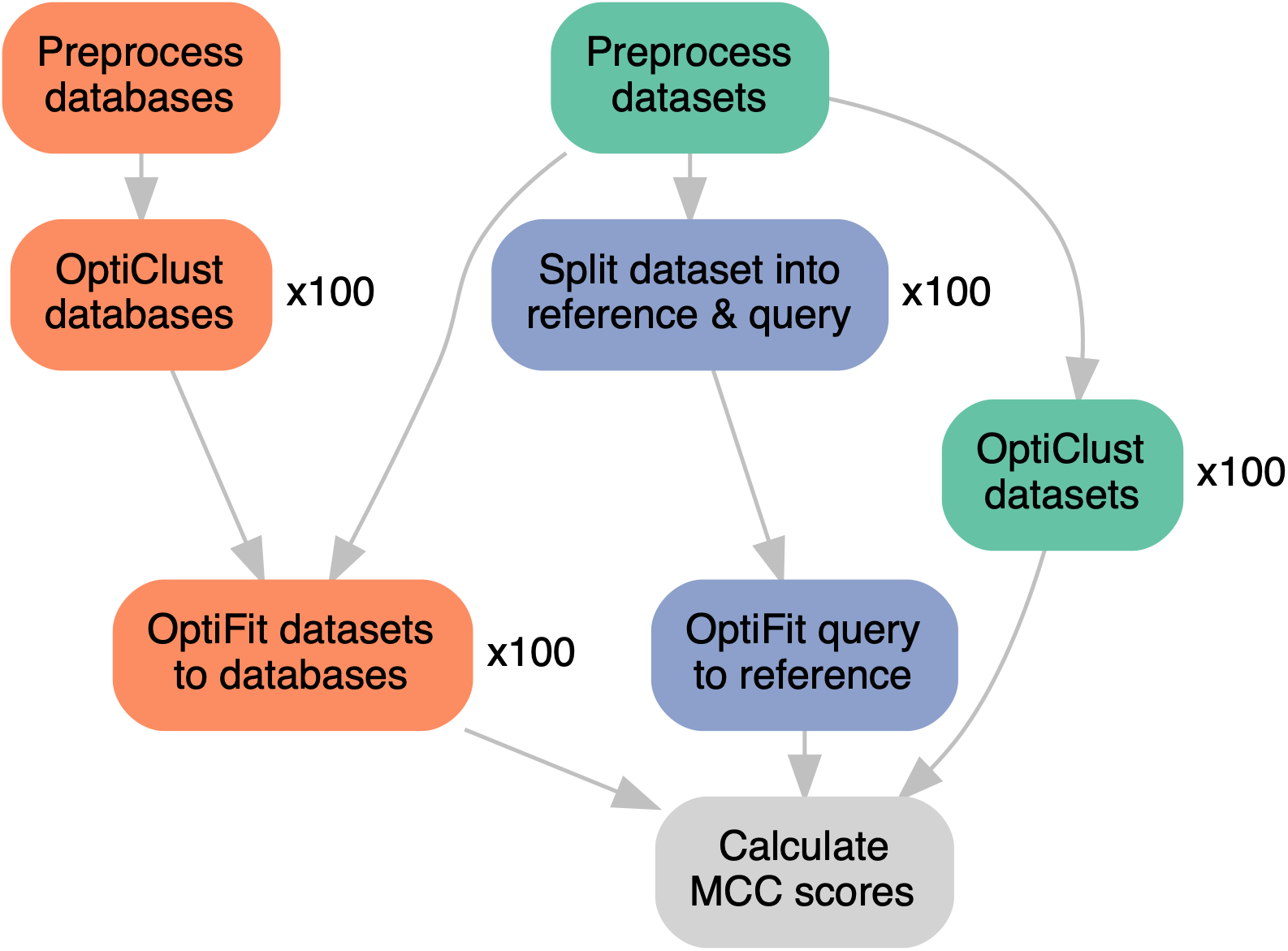
The Analysis Workflow. Reference sequences from Greengenes, the RDP, and SILVA were downloaded, preprocessed with mothur by trimming to the V4 region, and clustered *de novo* with OptiClust for 100 repetitions. Datasets from human, marine, mouse, and soil microbiomes were downloaded, preprocessed with mothur by aligning to the SILVA V4 reference alignment, then clustered *de novo* with OptiClust for 100 repetitions. Individual datasets were fit to reference databases with OptiFit; OptiFit was repeated 100 times for each dataset and database combination. Datasets were also randomly split into a reference and query fraction, and the query sequences were fit to the reference sequences with OptiFit for 100 repetitions. The final MCC score was reported for all OptiClust and OptiFit repetitions.

**Figure 3:**
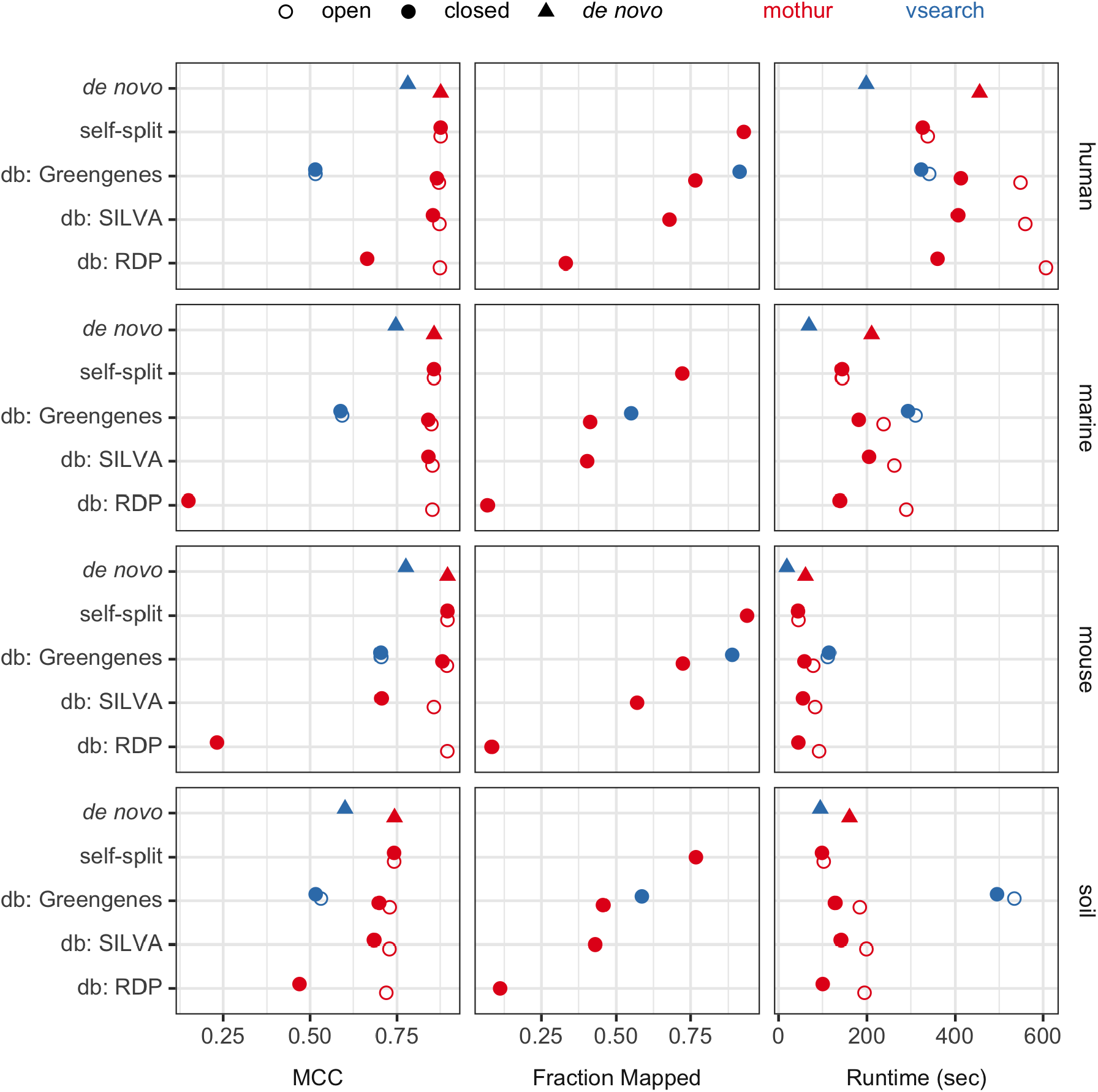
Benchmarking Results. The median MCC score, fraction of query sequences that mapped in closed-reference clustering, and runtime in seconds from repeating each clustering method 100 times. Each dataset underwent *de novo* clustering using OptiClust or reference-based clustering using OptiFit with one of two strategies; splitting the dataset and fitting 50% the sequences to the other 50%, or fitting the dataset to a reference database (Greengenes, SILVA, or RDP). Reference-based clustering was repeated with open and closed mode. For additional comparison, VSEARCH was used for *de novo* and reference-based clustering against the Greengenes database.

Since closed reference clustering does not cluster query sequences that could not be clustered into reference OTUs, an additional measure of clustering performance to consider is the fraction of query sequences that were able to be clustered. On average, more sequences were clustered with Greengenes as the reference (59.1%) than with SILVA (50.0%) or with the RDP (9.8%) (Figure 3). This mirrored the result reported above that Greengenes produced better OTUs in terms of MCC score than either SILVA or RDP. Note that *de novo* and open reference clustering methods always cluster 100% of sequences into OTUs. The database chosen affects the final closed reference OTU assignments considerably in terms of both MCC score and fraction of query sequences that could be clustered into the reference OTUs.

Despite the drawbacks, closed reference methods have been used when fast execution speed is required, such as when using very large datasets (14). To compare performance in terms of speed, we repeated each OptiFit and OptiClust run 100 times and measured the execution time. Across all dataset and database combinations, closed reference OptiFit outperformed both OptiClust and open reference OptiFit (Figure 3). For example, with the human dataset fit to SILVA reference OTUs, the average run times in seconds were 406.8 for closed reference OptiFit, 455.3 for *de novo* clustering the dataset, and 559.4 for open reference OptiFit. Thus, the OptiFit algorithm continues the precedent that closed reference clustering sacrifices OTU quality for execution speed.

To compare to the reference clustering methods used by QIIME2, we clustered each dataset with VSEARCH against the Greengenes database of OTUs previously clustered at 97% sequence similarity. Each reference OTU from the Greengenes 97% database contains one reference sequence, and VSEARCH maps sequences to the reference based on each individual query sequence’s similarity to the single reference sequence. In contrast, OptiFit accepts reference OTUs which each may contain multiple sequences, and the sequence similarity between all query and reference sequences is considered when assigning sequences to OTUs. In closed reference mode, OptiFit produced 27.2% higher quality OTUs than VSEARCH, but VSEARCH was able to cluster 24.8% more query sequences than OptiFit to the Greengenes reference database (Figure 3). This is because VSEARCH only considers the distances between each query sequence to the single reference sequence, while OptiFit considers the distances between all pairs of reference and query sequences in an OTU. When open reference clustering, OptiFit produced higher quality OTUs than VSEARCH against the Greengenes database, with median MCC scores of 0.86 and 0.56, respectively. In terms of run time, OptiFit outperformed VSEARCH in both closed and open reference mode by 54.6% and 49.5% on average, respectively. Thus, the more stringent OTU definition employed by OptiFit, which prefers the query sequence to be similar to all other sequences in the OTU rather than to only one sequence, resulted in fewer sequences being clustered to reference OTUs than when using VSEARCH, but caused OptiFit to outperform VSEARCH in terms of both OTU quality and execution time.

### Reference clustering with split datasets

When performing reference clustering against public databases, the database chosen greatly affects the quality of OTUs produced. OTU quality may be poor when the reference database consists of sequences that are too unrelated to the samples of interest, such as when samples contain novel populations. While *de novo* clustering overcomes the quality limitations of reference clustering to databases, OTU assignments are not consistent when new sequences are added. Researchers may wish to cluster new sequences to existing OTUs or to compare OTUs across studies. To determine how well OptiFit performs for clustering new sequences to existing OTUs, we employed a split dataset strategy, where each dataset was randomly split into a reference fraction and a query fraction. Reference sequences were clustered *de novo* with OptiClust, then query sequences were clustered to the *de novo* OTUs with OptiFit.

First, we tested whether OptiFit performed as well as *de novo* clustering when using the split dataset strategy with half of the sequences selected for the reference by a simple random sample (a 50% split) (Figure 3; self-split). OTU quality was similar to that from OptiClust regardless of mode (0.029% difference in median MCC). In closed reference mode, OptiFit was able to cluster 84.8% of query sequences to reference OTUs with the split strategy, a great improvement over the average 59.1% of sequences clustered to the Greengenes database. In terms of run time, closed and open reference OptiFit performed faster than OptiClust on whole datasets by 34.7% and 33.5%, respectively. The split dataset strategy also performed 13.5% faster than the database strategy in closed reference mode and 43.5% faster in open reference mode. Thus, reference clustering with the split dataset strategy creates as high quality OTUs as *de novo* clustering yet at a faster run time, and fits far more query sequences than the database strategy.

While we initially tested this strategy using a 50% split of the data into reference and query fractions, we next investigated whether there was an optimal reference fraction size. To identify the best reference size, reference sets with 10% to 90% of the sequences were created, with the remaining sequences used for the query (Figure 4). OTU quality was remarkably consistent across reference fraction sizes. For example, splitting the human dataset 100 times yielded a coefficient of variation (i.e. the standard deviation divided by the mean) of 0.00022 for the MCC score across all fractions. Run time generally decreased as the reference fraction increased; for the human dataset, the median run time was 364.1 seconds with 10% of sequences in the reference and 291.3 seconds with 90% of sequences in the reference. In closed reference mode, the fraction of sequences that mapped increased as the reference size increased; for the human dataset, the median fraction mapped was 0.85 with 10% of sequences in the reference and 0.95 with 90% of sequences in the reference. These trends held for the other datasets as well. Thus, the reference fraction did not affect OTU quality in terms of MCC score, but did affect the run time and the fraction of sequences that mapped during the closed reference clustering.

**Figure 4:**
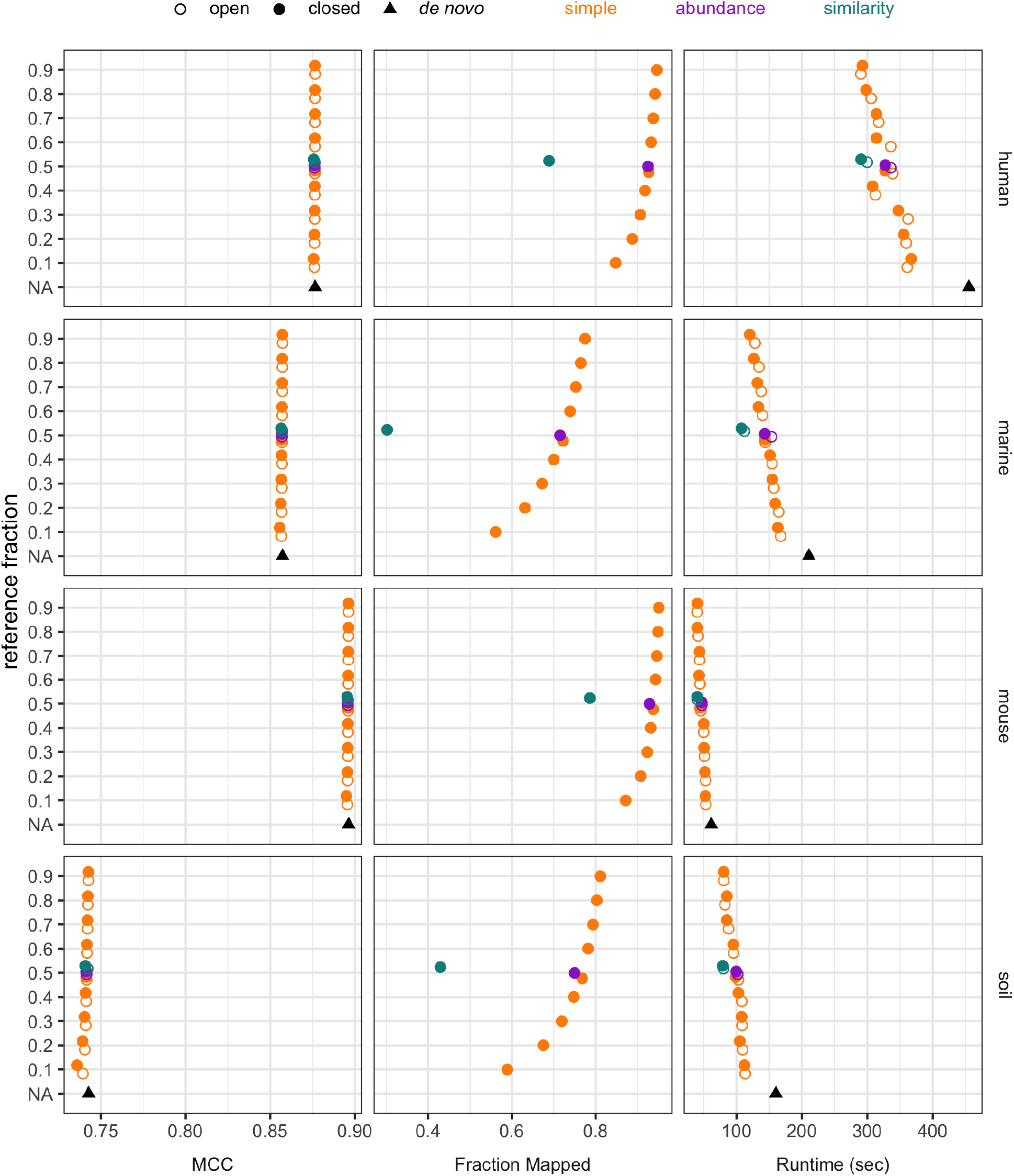
Split dataset strategy. The median MCC score, fraction of query sequences that mapped in closed-reference clustering, and runtime in seconds from repeating each clustering method 100 times. Each dataset was split into a reference and query fraction. Reference sequences were selected via a simple random sample, weighting sequences by relative abundance, or weighting by similarity to other sequences in the dataset. With the simple random sample method, dataset splitting was repeated with reference fractions ranging from 10% to 90% of the dataset and for 100 random seeds. *De novo* clustering each dataset is also shown for comparison.

After testing the split strategy using a simple random sample to select the reference sequences, we then investigated other methods of splitting the data. We tested three methods for selecting the fraction of sequences to be used as the reference at a size of 50%: a simple random sample, weighting sequences by relative abundance, and weighting by similarity to other sequences in the dataset (Figure 4). OTU quality in terms of MCC was similar across all three sampling methods (median MCC of 0.87). In closed-reference clustering mode, the fraction of sequences that mapped were similar for simple and abundance-weighted sampling (median fraction mapped of 0.85 and 0.84, respectively), but worse for similarity-weighted sampling (median fraction mapped of 0.56). While simple and abundance-weighted sampling produced better quality OTUs than similarity-weighted sampling, OptiFit performed faster on similarity-weighted samples with a median runtime of 93.8 seconds compared to 123.2 and 122.6 seconds for simple and abundance-weighted sampling, respectively. Thus, employing more complicated sampling strategies such as abundance-weighted and similarity-weighted sampling did not confer any advantages over selecting the reference via a simple random sample, and in fact decreased OTU quality in the case of similarity-weighted sampling.

## Discussion

We developed a new algorithm for clustering sequences to existing OTUs and have demonstrated its suitability for reference-based clustering. OptiFit makes the iterative method employed by OptiClust available for tasks where reference-based clustering is required. We have shown that OTU quality is similar between OptiClust and OptiFit in open reference mode, regardless of strategy employed. Open reference OptiFit performs slower than OptiClust due to the additional *de novo* clustering step, so users may prefer OptiClust for tasks that do not require reference OTUs.

When clustering to public databases, OTU quality dropped in closed reference mode to different degrees depending on the database and dataset source, and no more than half of query sequences were able to be clustered into OTUs across any dataset/database combination. This may reflect limitations of reference databases, which are unlikely to contain sequences from novel microbes. This drop in quality was most notable with the RDP reference, which contained only 16,192 sequences compared to 173,648 sequences in SILVA and 174,979 in Greengenes. Note that Greengenes has not been updated since 2013 at the time of this writing, while SILVA and the RDP are updated regularly. We recommend that users who require an independent reference database opt for large databases with regular updates and good coverage of microbial diversity for their environment. Since OptiClust still performs faster than open reference OptiFit and creates higher quality OTUs than closed reference OptiFit with the database strategy, we recommend using OptiClust rather than clustering to a database whenever consistent OTUs are not required.

The OptiClust and OptiFit algorithms produced higher quality OTUs than VSEARCH in open reference, closed reference, or *de novo* modes. However, VSEARCH was able to cluster more sequences to OTUs than OptiFit in closed reference mode. While both OptiFit and VSEARCH use a distance or similarity threshold for determining how to cluster sequences into OTUs, VSEARCH is more permissive than OptiFit regardless of mode. The OptiFit and OptiClust algorithms use all of the sequences to define an OTU, preferring that all pairs of sequences (including reference and query sequences) in an OTU are within the distance threshold in order to maximize the MCC. In contrast, VSEARCH only requires each query sequence to be similar to the single centroid sequence that seeded the OTU. Because of this, VSEARCH sacrifices OTU quality by allowing more dissimilar sequences to be clustered into OTUs.

When clustering with the split dataset strategy, OTU quality was remarkably similar when reference sequences were selected by a simple random sample or weighted by abundance, but quality was slightly worse when sequences were weighted by similarity. We recommend using a simple random sample since the more sophisticated reference selection methods do not offer any benefit. The similarity in OTU quality between OptiClust and OptiFit with this strategy demonstrates the suitability of using OptiFit to cluster sequences to existing OTUs, such as when comparing OTUs across studies. However, when consistent OTUs are not required, we recommend using OptiClust for *de novo* clustering over the split strategy with OptiFit since OptiClust is simpler to execute but performs similarly in terms of both run time and OTU quality.

Unlike existing reference-based methods that cluster query sequences to a single centroid sequence in each reference OTU, OptiFit considers all sequences in each reference OTU when clustering query sequences, resulting in OTUs of a similar high quality as those produced by the *de novo* OptiClust algorithm. Potential applications include clustering sequences to reference databases, comparing taxonomic composition of microbiomes across different studies, or using OTU-based machine learning models to make predictions on new data. OptiFit fills the missing option for clustering query sequences to existing OTUs that does not sacrifice OTU quality for consistency of OTU assignments.

## Materials and Methods

### Data Processing Steps

We downloaded 16S rRNA gene amplicon sequences from four published datasets isolated from soil (8), marine (9), mouse gut (10), and human gut (11) samples. These datasets contain sequences from the V4 region of the 16S rRNA gene and represent a selection of the broad types of natural communities that microbial ecologists study. We processed the raw sequences using mothur according to the Schloss Lab MiSeq SOP (15) and accompanying study by Kozich *et al*. (16). These steps included trimming and filtering for quality, aligning to the SILVA reference alignment (12), discarding sequences that aligned outside the V4 region, removing chimeric reads with UCHIME (17), and calculating distances between all pairs of sequences within each dataset prior to clustering.

### Reference database clustering

To generate reference OTUs from public databases, we downloaded sequences from the Greengenes database (v13_8_99) (6), SILVA non-redundant database (v132) (12), and the Ribosomal Database Project (v16) (13). These sequences were processed using the same steps outlined above followed by clustering sequences into *de novo* OTUs with OptiClust. Processed reads from each of the four datasets were clustered with OptiFit to the reference OTUs generated from each of the three databases. When reference clustering with VSEARCH, processed datasets were clustered directly to the unprocessed Greengenes 97% OTU reference alignment, since this method is how VSEARCH is typically used by the QIIME2 software for reference-based clustering (7, 18).

### Split dataset clustering

For each dataset, half of the sequences were selected to be clustered *de novo* into reference OTUs with OptiClust. We used three methods for selecting the subset of sequences to be used as the reference: a simple random sample, weighting sequences by relative abundance, and weighting by similarity to other sequences in the dataset. Dataset splitting was repeated with 100 random seeds. With the simple random sampling method, dataset splitting was also repeated with reference fractions ranging from 10% to 90% of the dataset. For each dataset split, the remaining query sequences were clustered into the reference OTUs with OptiFit.

### Benchmarking

OptiClust and OptiFit randomize the order of query sequences prior to clustering and employ a random number generator to break ties when OTU assignments are of equal quality. As a result, they produce slightly different OTU assignments when repeated with different random seeds. To capture any variation in OTU quality or execution time, clustering was repeated with 100 random seeds for each combination of parameters and input datasets. We used the benchmark feature provided by Snakemake to measure the run time of every clustering job. We calculated the MCC on each set of OTUs to quantify the quality of clustering, as described by Westcott *et al*. (1).

## Data and Code Availability

We implemented the analysis workflow in Snakemake (19) and wrote scripts in R (20), Python (21), and GNU bash (22). Software used includes mothur v1.47.0 (23), VSEARCH v2.15.2 (5), the tidyverse metapackage (24), R Markdown (25), ggraph (26), ggtext (27), numpy (28), the SRA toolkit (29), and conda (30). The complete workflow and supporting files required to reproduce this manuscript are available at https://github.com/SchlossLab/Sovacool_OptiFit_2021.

## Acknowledgements

We thank members of the Schloss Lab for their feedback on the figures.

KLS received support from the NIH Training Program in Bioinformatics (T32 GM070449). Salary support for PDS came from NIH grants R01CA215574 and U01AI124255. The funders had no role in study design, data collection and interpretation, or the decision to submit the work for publication.

## Author Contributions

KLS wrote the analysis code, evaluated the algorithm, and wrote the original draft of the manuscript. SLW designed and implemented the OptiFit algorithm and assisted in debugging the analysis code. MBM and GAD contributed analysis code. PDS conceived the study, supervised the project, and assisted in debugging the analysis code. All authors reviewed and edited the manuscript.

